# Divergent Biological Consequences of APOE Isoforms Across Industrialized and Non-Industrial Environments

**DOI:** 10.64898/2026.02.24.706846

**Authors:** Marina M. Watowich, Rachel M. Petersen, Layla Brassington, Audrey M. Arner, Grace Rodenberg, Tan Bee Ting A/P Tan Boon Huat, Kar Lye Tam, Em Schellenberg, Izandis bin Mohd Sayed, Echwa John, John C. Kahumbu, Benjamin Muhoya, Michael Gurven, Benjamin C. Trumble, Sospeter Ngoci Njeru, Dino Martins, Julien F. Ayroles, Yvonne A. L. Lim, Vivek V. Venkataraman, Ian J. Wallace, Thomas S. Kraft, Amanda J. Lea

## Abstract

The apolipoprotein ε4 (APOE ε4) isoform directly alters cholesterol and immune biology and is associated with an increased risk of neurodegenerative and cardiometabolic disease in industrialized settings; nevertheless, APOE ε4—which is ancestral in humans—has persisted over evolutionary time. One potential explanation is that the costs and benefits of APOE ε4 were significantly different in the environments in which humans evolved compared to those we experience today. In support, previous work has suggested that living in a high pathogen environment, engaging in high levels of physical activity, or eating a low fat diet can dampen the detrimental effects of APOE ε4, and has revealed positive effects for fertility. However, direct tests of whether APOE isoforms are associated with different biological outcomes in non-industrial versus industrialized contexts are lacking. Working with the Turkana of Kenya and the Orang Asli of Peninsular Malaysia—two Indigenous groups in which individuals of shared ancestry span a continuum of subsistence, non-industrial to urban, industrialized lifestyles—we investigated how *APOE* genotypes impact cholesterol, immunological, and reproductive traits and tested for genotype x environment (GxE) interactions. First, we confirmed established genotype effects across lifestyles, showing that more APOE ε4 alleles are associated with higher total cholesterol, higher LDL cholesterol, and lower HDL cholesterol. Second, we tested for lifestyle interactions, finding lifestyle-dependent effects of genotype on innate immune biomarkers in the Orang Asli but not Turkana. Finally, we show that more APOE ε4 alleles are correlated with an extended reproductive lifespan, however this effect is relatively weak, is not consistent across populations, and does not correspond with a higher reproductive output. Together, our study provides evidence that industrialized environments can modify the biology of APOE ε4; however, we find that APOE ε4 is not universally beneficial in non-industrial contexts, highlighting the role of local environmental variation in determining its specific costs and benefits.

## Introduction

Humans maintain 3 isoforms of the Apolipoprotein E protein—APOE ε4, APOE ε3, and APOE ε2—which each confer distinct and pleiotropic effects on cholesterol and innate immune traits [1,2]. APOE ε4 is well known as a major disease risk marker: in urban, industrialized settings, individuals homozygous for the ε4 isoform have a ∼12-fold higher risk of Alzheimer’s disease, a ∼1.5-fold higher risk of cardiovascular diseases (CVD), and an overall higher risk of premature death [3–5]. However, the APOE ε4 allele is considered ancestral in humans and shows minimal evidence of being selected against, raising the question of why it persists despite clear costs [6,7].

Two evolutionary theories of aging may explain the persistence of APOE ε4: first, that APOE ε4’s harmful effects are sufficiently concentrated at older ages in the “selection shadow” (consistent with mutation accumulation theory), and second, that APOE ε4 may provide benefits during reproductive years that outweigh its later life costs (consistent with antagonistic pleiotropy theory) [8–10]. A third, non-mutually exclusive hypothesis is that the costs and benefits of APOE ε4 are significantly different in modern, industrialized environments relative to those in which humans evolved, hindering our appraisal of its adaptive value [11–14]. In support of this third hypothesis, recent work has suggested that APOE ε4 can be less detrimental under preindustrial conditions: for example, in the presence of high pathogen loads, high physical activity levels, or a non-processed, anti-inflammatory diet, APOE ε4 has weaker effects on risk of both CVD and cognitive impairment [15–21]. Additionally, work with Tsimane forager-horticulturalists in Bolivia suggests that APOE ε4 may even provide benefits to fertility and cognition in non-industrial, energy-limited, high pathogen settings, contrary to what has been observed in industrialized environments [16,22–25]. Although recent work across populations (e.g., studies of the Tsimane versus the US) points toward greater benefits and dampened costs of APOE ε4 in non-industrial relative to industrialized environments, more direct tests for such genotype x environment interactions within populations are generally lacking.

The APOE protein is predominately produced in the liver, but the gene is also expressed by immune (e.g., macrophages) and neuronal (e.g., microglia, astrocytes) cell types [26]. The primary role of APOE is as a lipid transporter, binding to cholesterol and other phospholipids and delivering them throughout the body. Each APOE isoform has distinct cholesterol affinities: APOE ε2 and to a lesser extent APOE ε3 preferentially bind to cholesterol-rich, high-density lipoproteins (HDL), whereas APOE ε4 preferentially binds to triglyceride-rich, very low-density lipoproteins (VLDL) [27]. APOE ε4 also exhibits faster turnover of VLDL to LDL, but less effective clearance of cholesterol from circulation, resulting in higher amounts of total cholesterol and LDL in circulation for APOE ε4 as compared to APOE ε3 and especially APOE ε2 carriers [13,28,29]. Each APOE isoform also has distinct effects on immune function: APOE ε4 has been associated with baseline downregulation of the innate immune system in industrialized settings (e.g., lower CRP, IL6, and TNF-α levels) [30,31]) but stronger activation when engaged [32]. Further, relative to APOE ε2 and APOE ε3, the APOE ε4 isoform confers decreased risk of giardia, cryptosporidium, and hepatitis B and C, but increased susceptibility to HSV-1, *C. pneumoniae*, and HIV [33–35].

Several ideas have been proposed to explain why APOE ε4-associated cholesterol and/or immune traits may have been beneficial throughout human evolution. First, increased cholesterol availability could have been helpful in metabolically demanding thermal environments, such as high altitudes or latitudes, though evidence for this idea is mixed [36]. Second and more supported, APOE ε4 may provide several important immune benefits that are obscured or perturbed in industrialized environments: 1) cholesterol is both a component of immune cell membranes and a modulator of inflammatory signaling, such that increased circulating cholesterol could fuel immune responses in energy-limited, non-industrial environments [15,25,37,38]; and 2) the rapid and intense immune engagement characteristic of APOE ε4 carriers may be beneficial in high pathogen environments, but detrimental in industrialized environments if chronic stimulation is also induced by obesity, diet, or other sterile sources of damage-associated molecular patterns [15,39–43]. Finally, some combination of APOE ε4 carrier’s cholesterol and immune traits may create a fertility advantage in non-industrial settings: though the specific mechanism is unknown, APOE ε4 has been associated with earlier reproduction, shorter interbirth intervals, and greater lifetime reproductive success in energy-limited, high-pathogen settings in Bolivia, Ghana, and Ecuador [22–24]. Importantly, these same fertility benefits have not been observed in similar-sized studies of industrialized cohorts (reviewed in [22]). Taken together, these findings highlight the need to understand how APOE isoforms impact cholesterol, immune function, and fertility in non-industrial environments relevant to human evolution, and further, to directly test if these relationships are altered by industrialized environments.

By working with two non-industrial populations that are experiencing rapid but heterogeneous lifestyle change, our goal is to test how exposure to industrialized, urban lifestyles alter the biological consequences of APOE isoforms within a common genetic background [44,45].

Specifically, we worked with Turkana pastoralists of Northern Kenya and Orang Asli hunter-gatherers and horticulturalists of Peninsular Malaysia, two Indigenous groups living across a recently formed gradient of non-industrial to industrialized environments [46–48]. As part of long-term anthropological and biomedical work with each group, we collected data on APOE genotypes, cholesterol (total, LDL, and HDL levels), innate immune (5 part white blood cell differentials, CRP and cytokine levels, blood transcriptomes), and fertility traits (age at menarche, age at first reproduction, age-specific fertility, completed fertility, age at menopause) for 1965 individuals (1104 Turkana and 861 Orang Asli). These data were collected across a comparable, continuous, well-characterized gradient from remote, small-scale, subsistence-based communities to highly urban, industrialized, and market-integrated locations [47]. Using parallel datasets from both groups, we tested the predictions that: 1) regardless of lifestyle, APOE ε4 would predict higher total and LDL cholesterol levels, but that 2) industrialized lifestyles might alter APOE genotype-innate immune trait associations. Finally, we tested for 3) lifestyle-dependent effects of APOE genotype on female fertility outcomes, with the prediction that APOE ε4 carriers would experience greater benefits in non-industrial environments (Figure 1). Together, these analyses provide a rare opportunity to test for GxE interactions within two independent groups spanning an industrialized lifestyle gradient, allowing us to evaluate the evolutionary significance of a major disease-associated genetic locus.

**Figure 1.**
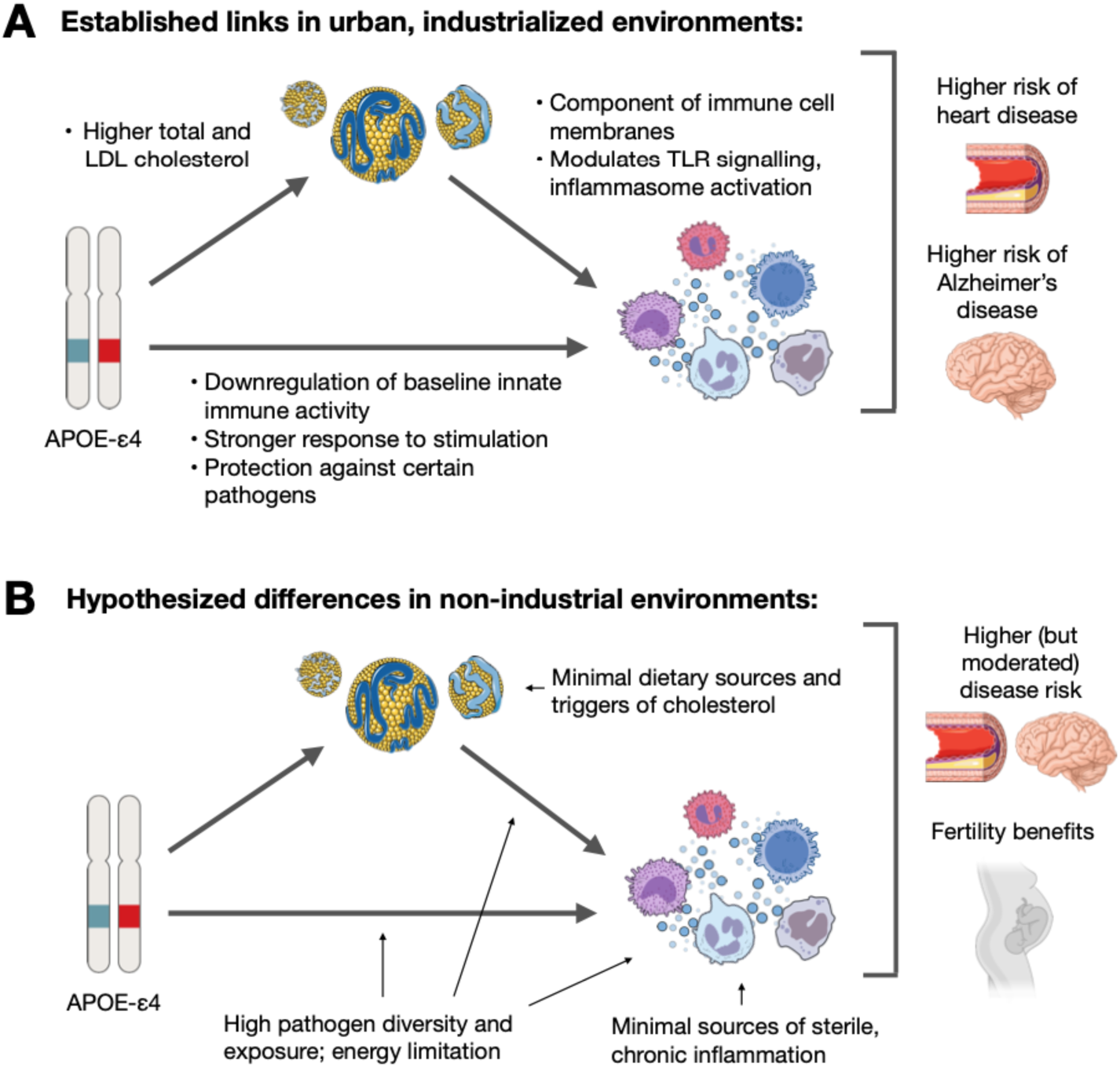
Conceptual model and hypothesized differences in biological relationships between non-industrial versus industrialized environments. (A) Previous work has established that in urban, industrialized settings, 1) APOE ε4 carriers exhibit elevated total and LDL cholesterol; 2) APOE ε4 modulates innate immune function; and 3) cholesterol availability (driven by APOE genotype and other sources) impacts the formation of immune cell membranes, Toll-like receptor (TLR) signaling, and inflammasome activation. (B) In non-industrial environments characterized by limited dietary lipid availability, high pathogen burden, high physical activity, and constrained energetic resources, APOE ε4 may have different biological correlates that dampen its effects on disease [16,49]. We hypothesize that, across genotypes, non-industrial lifestyles will lower total and LDL cholesterol and generate greater immune activation from non-sterile sources. Further, we expect that non-industrial lifestyles will reveal different APOE ε4-immune and cholesterol-immune relationships relative to industrialized contexts, because both cholesterol accumulation and the reactive, APOE ε4 innate immune phenotype may be advantageous in the absence of lifestyle-induced hypercholesterolemia or sterile, chronic inflammation. Adapted from [15]. Illustrations created with resources from NIH BioArt and Servier Medical Art, licensed under CC BY 4.0.

## Results

### Lifestyle variation provides opportunities to test for genotype x environment interactions

We drew on questionnaire, immune, cholesterol, and fertility data from two long-term projects: the Turkana Health and Genomics Project (THGP) and the Orang Asli Health and Lifeways Project (OA HeLP) [46,50] (Figure 2A and Figure S1). Both projects work with communities that span a gradient from non-industrial, remote-living, subsistence-based to urban, market-integrated, industrialized environments. Briefly, the Turkana are traditionally nomadic pastoralists living in arid regions of Northwest Kenya, but increasing droughts, expansion of markets, and changing economy in Kenya has led to an increased reliance on wage labor and urban infrastructure in recent decades. The term “Orang Asli” refers to 19 ethnolinguistic groups from three major subgroups (Senoi, Negrito, Proto Malay) that together comprise the Indigenous peoples of Peninsular Malaysia. Orang Asli have traditionally practiced a variety of subsistence strategies including hunting, gathering, horticulture, and fishing, but due especially to ongoing deforestation and government-led resettlement initiatives, many Orang Asli are now highly market-integrated and more focused on wage labor.

**Figure 2.**
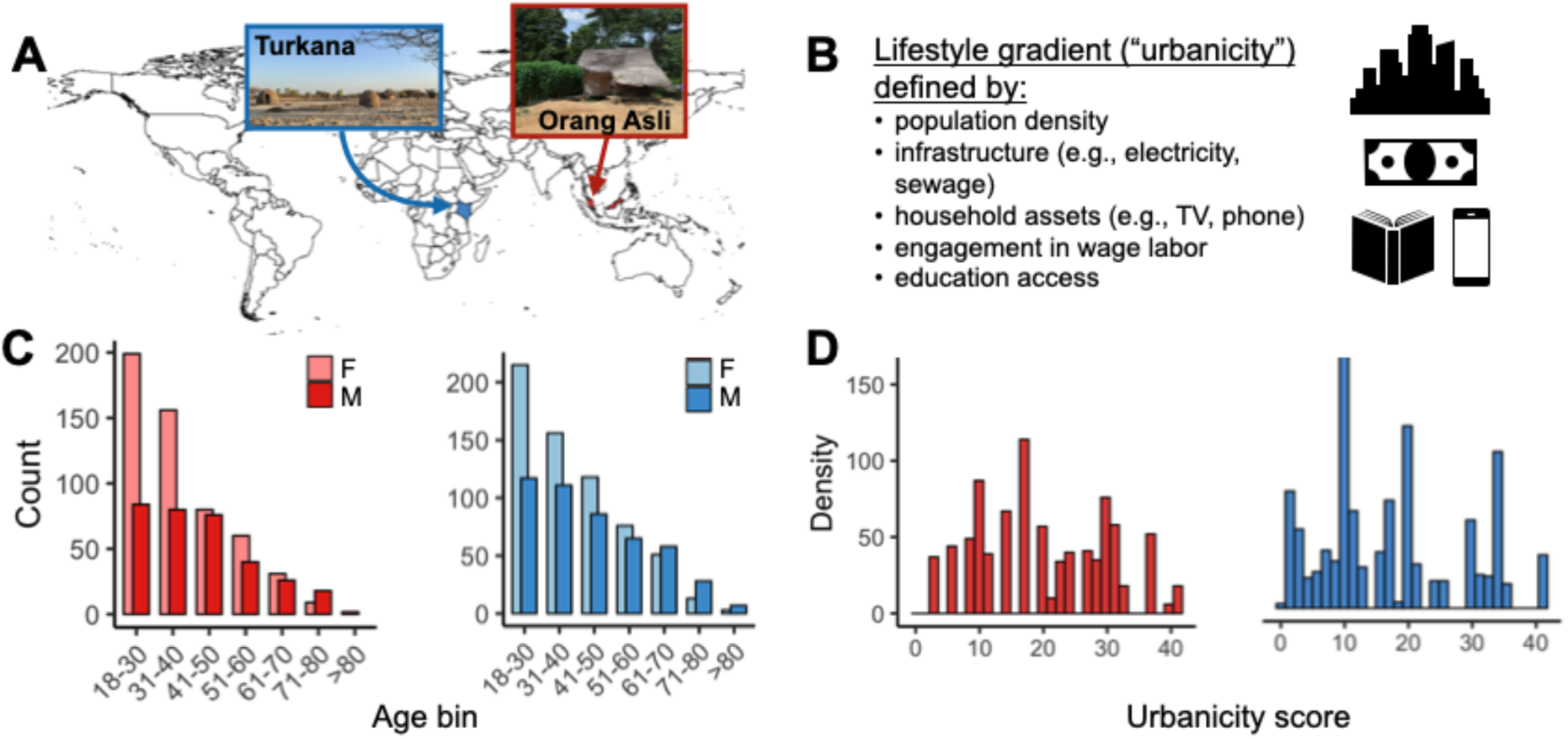
Study populations and lifestyle variation. (A) World map highlighting the countries where the Turkana and Orang Asli live (note Orang Asli reside specifically in Peninsular Malaysia). Pictures (which were taken by the authors) show traditional housing in non-industrial settings. (B) Description of a previously validated “urbanicity” score used to quantify the continuum of non-industrial to industrialized lifestyles in a comparable way across both groups. (C) Sex and age distributions of study participants (F=female; M=male). (D) Range of urbanicity scores for the Turkana and Orang Asli sample sets. Across all panels, data from Turkana are shown in blue and data from Orang Asli are in red.

To operationalize this lifestyle variation, our group has previously adapted an existing measure of “urbanicity” [51] that sums information about population density, average access to infrastructure (electricity, sewage), ownership of market-derived good (televisions, mobile phones), subsistence activities, and education access within a given location [47] (Figure 2B, urbanicity scale construction is detailed in Supplementary Methods). While we refer to this measure as an “urbanicity” index (because this is the term used in the original publication [51]) we emphasize that it is meant to capture how industrialized and market-integrated each individual’s lifestyle is—which in our sample includes fully non-industrial lifestyles—and is not equivalent to rural-urban gradients in high income countries. In total, this study included 1104 Turkana participants from 67 locations (57% female, 43% male) and 861 Orang Asli participants from 30 locations (62% female, 38% male; Figure 2C and Table S1-2). These locations spanned a similar, wide range of urbanicity values in Kenya and Malaysia (Figure 2D).

Importantly, we have previously shown that urbanicity correlates with relevant aspects of energy balance (Figure 1): individuals living in locations of higher urbanicity have significantly lower physical activity levels in Orang Asli as well as greater access to processed foods, higher body fat percentages, and higher body mass indices in both groups [47,52]. We also confirmed these same correlations among APOE genotyped individuals in this study (linear model, all p < 0.05; Table S3). Further information on the THGP, OA HeLP, and participant communities are provided in the Supplementary Methods.

### APOE genotypes are generally in Hardy-Weinberg equilibrium

The three APOE isoforms are distinguished by just two amino acid substitutions (ε2: Cys112/Cys158; ε3: Cys112/Arg158; ε4: Arg112/Arg158). We inferred these substitutions by genotyping the rs429358 and rs7412 single nucleotide polymorphisms (SNPs), and found that APOE isoform frequencies were consistent with previous reports [53,54]: 46.4% of Turkana (n = 513/1104) and 35.4% of Orang Asli (n = 305/861) participants had at least one copy of APOE ε4, with only 8.1% of Turkana (n = 90/1105) and 4.1% of Orang Asli (n = 35/861) being ε4/ε4 homozygous (Figure 3A). Despite its associations with cardiovascular and neurodegenerative disease in industrialized societies, the frequency of APOE ε4 was more than three times higher than that of APOE ε2 in Turkana (allele frequency ε4 = 29.8%, ε2 = 9.4%) and 1.9 times higher in Orang Asli (allele frequency ε4 = 22.1%, ε2 = 11.1%). Both SNPs were in Hardy-Weinberg Equilibrium (HWE) in the Turkana (Haldane Exact Tests; rs429358, p = 0.25 and rs7412, p = 0.15). In the Orang Asli, the rs429358 SNP was in HWE, but the rs7412 was not (p = 0.008).

**Figure 3.**
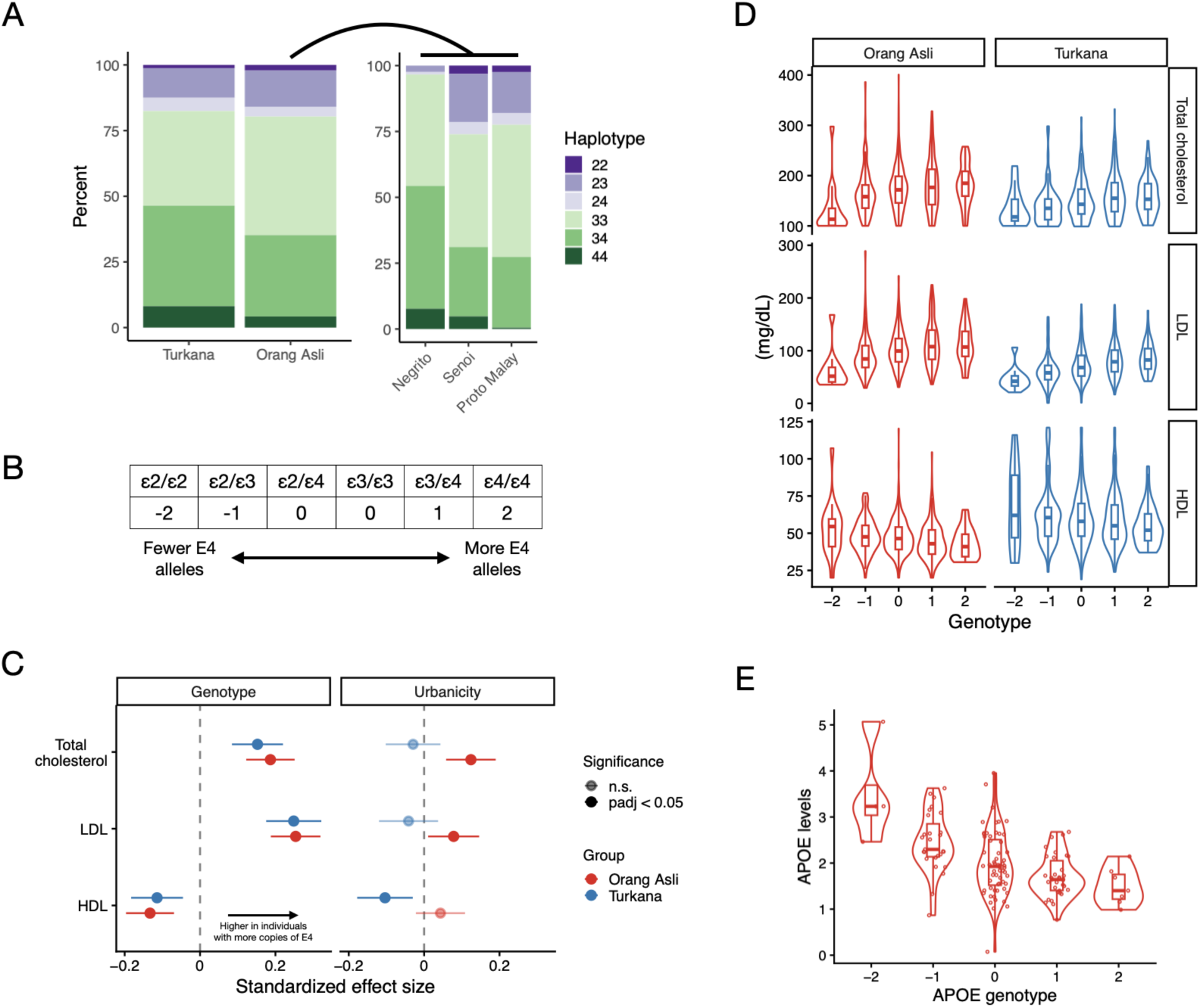
APOE ε4 is prevalent at moderate frequencies and is associated with higher blood cholesterol in Turkana and Orang Asli. (A) Frequencies of each apolipoprotein (APOE) haplotype in the Turkana and Orang Asli, with frequencies for major Orang Asli subgroups shown to the right. (B) The numeric values assigned to each APOE haplotype. (C) Forest plot of the effect of APOE (as a linear variable) on cholesterol measures from blood samples. Points represent the standardized effect size and bars represent 95% confidence intervals. Positive values indicate positive associations between cholesterol phenotypes and the linear genotype scale (i.e., individuals with more ε4 alleles) or living in higher urbanicity settings. (D) Cholesterol values per genotype category. Values shown are predicted from linear models of cholesterol controlling for age, sex, and urbanicity score. (E) Apolipoprotein E serum protein levels across APOE haplotypes in the Orang Asli. Values shown are predicted from linear models of protein levels controlling for age, sex, genotype, and urbanicity score.

However, because there is some known, slight genetic differentiation between the three Orang Asli subgroups [55], we also repeated our analyses following self-reported ancestry stratification and found that almost all subgroup-specific tests supported HWE, with the exception of rs429358 in Proto Malay (p = 0.03; Table S4).

### APOE ε4 is associated with higher total and LDL cholesterol levels, and lower HDL cholesterol levels, across the industrialization gradient

Based on previous studies, we expected APOE ε4 to correlate with higher total and LDL cholesterol levels across groups and environments and found that this was the case: Turkana and Orang Asli with more APOE ε4 copies had higher total cholesterol (linear model, Turkana p_FDR_ = 2.82 x 10^-3^, Orang Asli p_FDR_ = 1.25 x 10^-3^), higher LDL cholesterol (Turkana p_FDR_ = 1.57 x 10^-6^, Orang Asli p_FDR_ = 1.57 x 10^-6^), and lower HDL cholesterol levels (Turkana p_FDR_ = 4.92 x 10^-3^, Orang Asli p_FDR_ = 1.04 x 10^-3^; Figure S2 and Table S5). A somewhat unique feature of our study is that all 3 isoforms are represented in both groups, allowing us to compare both the APOE ε4 as well as the APOE ε2 isoform to the more common APOE ε3 isoform. Consistent with previous work, we found that individuals with more ε2 alleles had lower total cholesterol (Turkana p_FDR_ = 9.22 x 10^-5^, Orang Asli p_FDR_ = 6.92 x 10^-8^), lower LDL cholesterol (Turkana p_FDR_ = 1.53 x 10^-7^, Orang Asli p_FDR_ = 2.01 x 10^-10^), and higher HDL cholesterol levels (Turkana p_FDR_ = 3.20 x 10^-2^, Orang Asli p_FDR_ = 5.01 x 10^-3^; Figure S2 and Table S5) relative to the other isoforms.

Because the APOE ε4 and APOE ε2 isoforms had essentially mirror opposite effects, we focused on modeling APOE genotypes as a continuous, linear variable for all remaining analyses. Specifically, we centered this measure on the APOE ε3/ε3 genotype, adding one per each APOE ε4 copy and subtracting one per each APOE ε2 copy (Figure 3B). We use this approach for our main text analyses, but present parallel results using the number of APOE ε4 or APOE ε2 copies as the predictor in the Supplementary Materials.

Using the continuous approach, we confirmed our expectation that genotype linearly impacts all three cholesterol traits in both groups (all p_FDR_ < 0.05; Figure 3C-D), as well as APOE serum protein levels measured for 132 Orang Asli participants (linear model, p = 2.49 x 10^-9^; Figure 3E). As expected, these effects are not lifestyle-dependent (all p_FDR_ for a genotype x lifestyle interaction > 0.1; Table S6-7). Nevertheless, greater urbanicity was associated with higher total (p_FDR_ = 4.43 x 10^-3^) and LDL cholesterol levels (p_FDR_ = 0.07, p_unadjusstested_ = 0.04), but not HDL cholesterol levels (p_FDR_ = 0.76), in Orang Asli (Figure 3C). In contrast, in Turkana, greater urbanicity was associated with lower HDL cholesterol levels (p_FDR_ = 1.85 x 10^-2^), but had no effects on total and LDL cholesterol levels (Figure 3C and Table S6-7). These results are notable because previous work has largely failed to identify urbanicity effects on lipid traits in either population [46,47,56], which are here identified when controlling for APOE genotype.

### APOE isoforms have lifestyle-dependent effects on the innate immune system

Because increasingly industrialized lifestyles are associated with modifications to energy balance, physical activity, and diet in our participant populations, as well as potentially pathogen exposure, we hypothesized that lifestyle should modify APOE genotype-innate immune trait associations (Figure 1). We tested this by assessing genotype x urbanicity interactions for a suite of immune biomarkers: innate immune cell counts (from a 5-part white blood cell differential), C-reactive protein (CRP) levels, plasma cytokine levels (Orang Asli only), and peripheral blood mononuclear (PBMC) gene expression levels (Orang Asli only). Several biomarkers exhibited evidence only for main effects of genotype, which generally pointed to innate immune downregulation among APOE ε4 carriers, consistent with previous work in industrialized contexts (Figure 4A, Figure S3, and Table S8-11). Specifically, in Orang Asli, continuous genotype trended toward a negative association with CRP levels (linear model, p_unadjusted_ = 0.01, p_FDR_ = 0.14; Figure 4B), indicating lower inflammation with more APOE ε4 alleles. Similarly, in our Orang Asli PBMC gene expression dataset (n = 9993 expressed genes, n samples = 745), we observed three genes significantly associated with APOE genotype at a false discovery rate of 10% (*LAMA5*, *PAIP2B*, *SIX5*). These genes are implicated in tissue development and integrity and each had higher relative expression in APOE ε4 carriers. We then performed gene set enrichment analyses after ranking genes based on their genotype effect sizes, and found enrichment of pathways associated with inflammatory and innate immune activity (e.g., p_FDR_ = 1.20 x 10^-13^ for “TNFα signaling via NFKB”, p_FDR_ = 9.73 x 10^-6^ for “interferon gamma response”, p_FDR_ = 1.47 x 10^-6^ for “inflammatory response”; Figure S4). The direction of these effects suggests that individuals with more APOE ε4 copies downregulate these processes relative to APOE ε3 and especially APOE ε2 carriers (Figure 4C-D).

**Figure 4.**
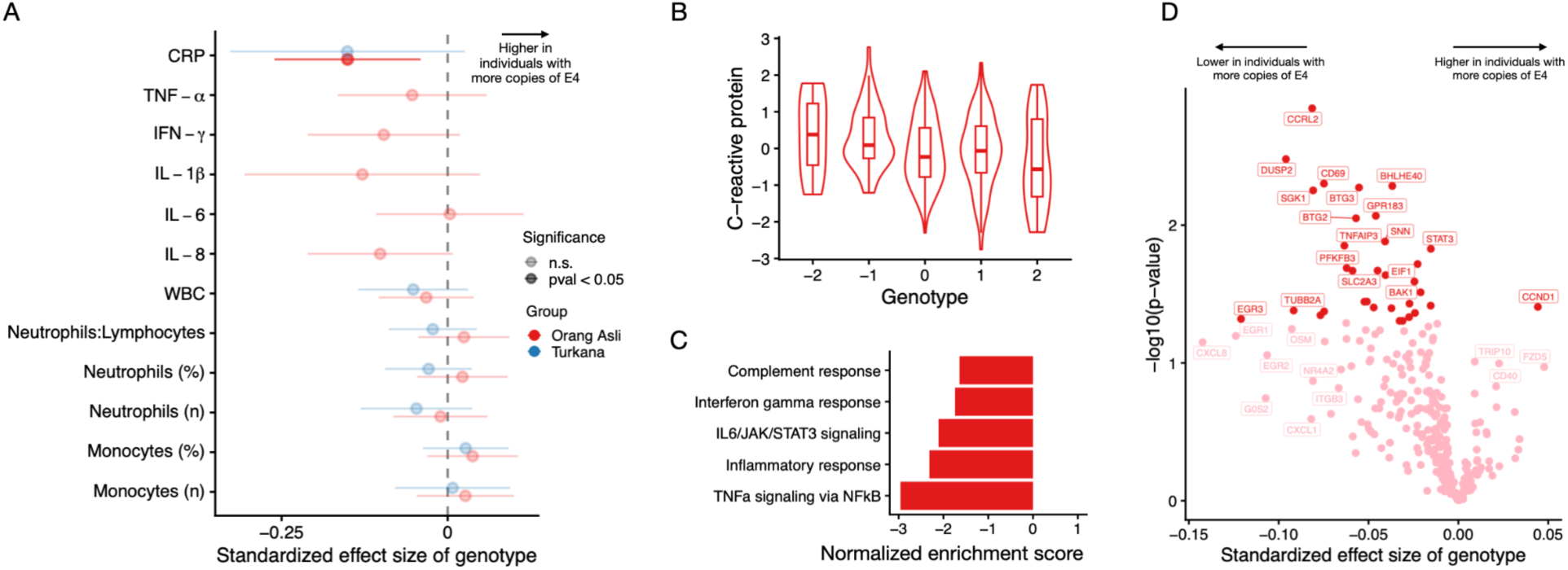
APOE ε4 is associated with lower inflammation and innate immune activity. (A) Forest plots of the effect of APOE genotype on innate immune cell count and percent, total WBC count, CRP, and innate immune biomarkers for Orang Asli (red) and Turkana (blue). (B) CRP values per genotype category predicted from linear models controlling for age and sex for Orang Asli. (C) Under-enrichment of GSEA hallmark terms related to inflammation and innate immune activity in gene expression from unstimulated PBMCs in Orang Asli. (D) Example volcano plot of genes in the TNF-ɑ signaling via NFκB pathway.

In Orang Asli, but not Turkana, several innate immune biomarkers trended toward evidence for GxE interactions (Figure 5A). Specifically, genotype effects on the percentage (p_FDR_ = 0.1, p_unadjusted_ = 1.48 x 10^-2^) and total abundance of neutrophils (p_FDR_ = 0.09, p_unadjusted_ = 6.41 x 10^-3^) in blood, and the total amount of white blood cells (p_FDR_ = 0.12, p_unadjusted_ = 2.75 x 10^-2^) trended toward depending on lifestyle. These interactions consistently manifested as more copies of APOE ε4 being associated with higher biomarker values in rural, non-industrial contexts, but lower biomarker values in urban, industrialized contexts (Figure 5B, Table S8-11). Thus, across the Orang Asli industrialization gradient, our results suggest that the previously reported immunosuppressive effects of APOE ε4 are magnified in more urban, industrialized contexts in which they have been overwhelmingly studied.

**Figure 5.**
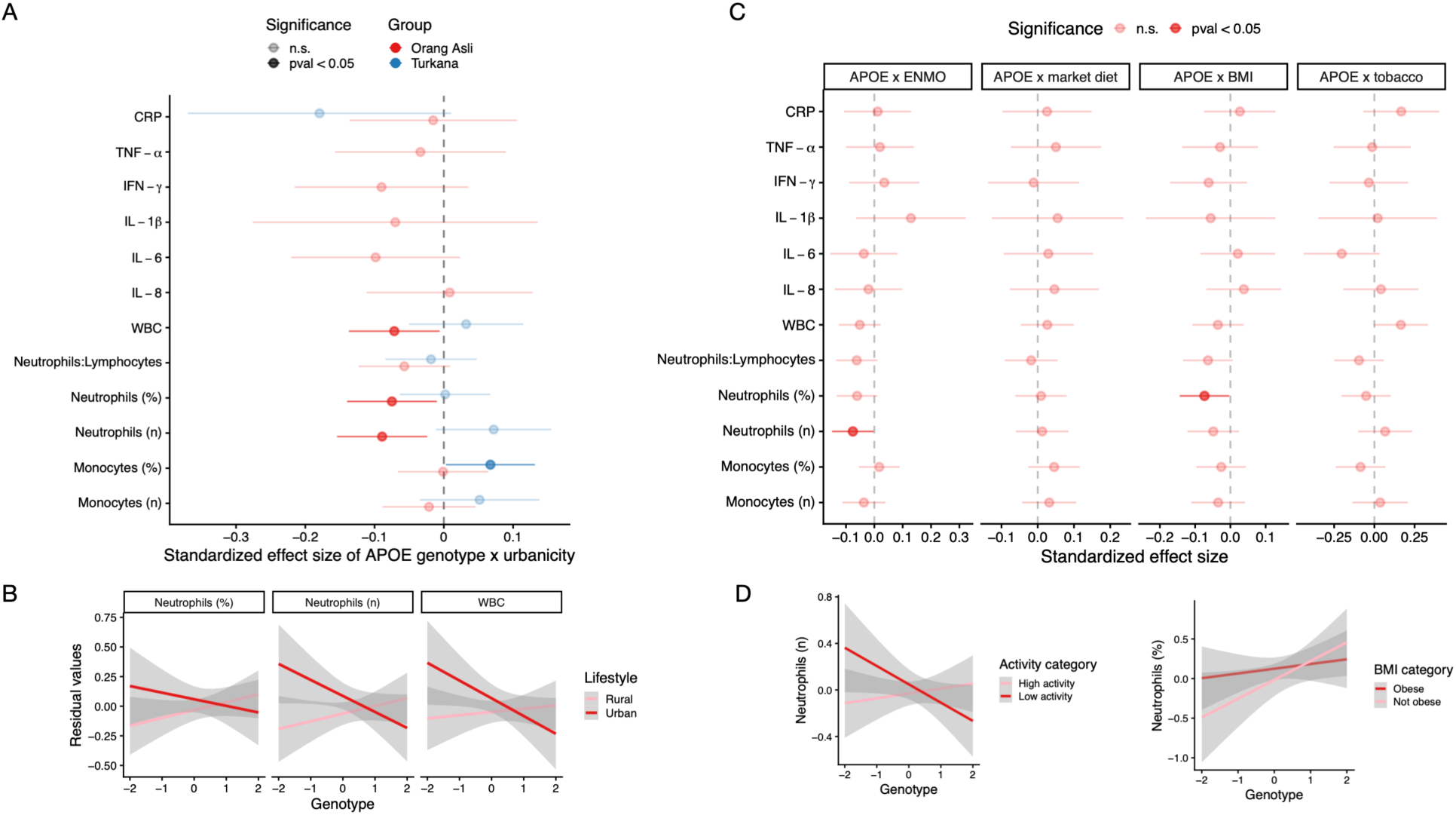
Many innate immune phenotypes exhibit environment x APOE genotype effects and the urbanicity index explains more variance than individual components of the urban environment. (A) Forest plot of the effect of APOE genotype x urbanicity effects on innate immune cell count and percent, total WBC count, CRP, and innate immune biomarkers for Orang Asli (red) and Turkana (blue). (B) Innate immune and inflammatory biomarker values across genotype for biomarkers showing genotype by environment interactive effects in Orang Asli. Values are predicted from linear models controlling for age, sex, APOE genotype, and urbanicity. (C) Forest plots of genotype by environment effects of several proximate, individual components of the urban environment on innate immune biomarkers (ENMO indicates physical activity, with Euclidean Norm Minus One or ENMO from accelerometry). (D) Innate immune biomarker values across genotype for biomarkers showing trending genotype by mean activity level and anthropometric nutritional status effects in Orang Asli. Values are predicted from linear models controlling for age, sex, APOE genotype, and ENMO or BMI.

To determine which facets of urban, industrialized environments could contribute to the GxE interactions observed in the Orang Asli, we also reran these analyses swapping out our urbanicity score for 1) body mass index (BMI), 2) an index of how much access each individual has to market-derived, processed foods [47], 3) usage of tobacco products (never versus sometimes or daily), and 4) objectively measured physical activity levels (measured as mean Euclidean Norm Minus One or ENMO from accelerometry) [52]. Each of these factors are components of the broader non-industrial to industrialized lifestyle continuum, are thought to perturb the innate immune system via sterile, chronic inflammation [40], and have been previously hypothesized as modifiers of APOE ε4’s effects [15,17,19–21]. None of these individual factors recapitulated GxE interactions as strongly as what we observed using our composite urbanicity score (Figure 5C). However, BMI came the closest, with trends toward GxE interactive effects on neutrophil percentage (p_FDR_ = 0.47, p_unadjusted_ = 4.01 x 10^-2^), and neutrophil to lymphocyte ratios (p_FDR_ = 0.47, p_unadjusted_ = 7.8 x 10^-2^; Figure 5D and Table S12).

Finally, for comparison to previous work in the Tsimane, we also tested whether lifestyle modified links between cholesterol availability and innate immune system function (Figure 1) [15]. We found that cholesterol-immune biomarker links varied across the industrialization gradient in heterogeneous ways, but with some consistent patterns. For example, in both populations, higher total cholesterol levels were associated with lower numbers of monocytes in non-industrial environments, but higher numbers of monocytes in urban environments (Figure S5 and Table S13-14). The patterns we observed in non-industrial individuals are consistent with previous findings in the Tsimane, and were interpreted as evidence that, in energy-limited contexts, high-cholesterol individuals are better able to tolerate infection [15].

### APOE genotype has weak and heterogeneous effects on reproductive traits

Finally, we tested whether APOE genotype impacts female reproductive traits and whether these effects vary across the industrialization gradient (Figure 6A). We focused on age at first birth (Turkana only), age at menarche (Turkana only), age-adjusted fertility, completed fertility, and age at menopause (Orang Asli only) (Table S15-16). All reproductive information was self-reported. Overall, we found minimal evidence for genotype or GxE interaction effects, but with a few population-specific results passing our significance thresholds. First, in Turkana, we found that genotype impacted age at menarche, with more APOE ε4 alleles correlating with earlier reproductive maturation (Figure 6B; n = 474, p = 0.048). Second, in Orang Asli, we found GxE interaction effects for age at menopause: APOE ε2 carriers experience an earlier age of menopause in rural compared to urban contexts, whereas APOE ε4 carriers experienced a more similar age at menopause regardless of urbanicity (Figure 6C; n = 454, p = 0.006). The weak, varied effects of APOE genotype on fertility outcomes that we observed were generally robust when excluding women currently on contraception (24.4% of women in OA and 18.9% of women in Turkana; Table S15-16) and to using different age cutoffs to define completed fertility (Figure S6).

**Figure 6.**
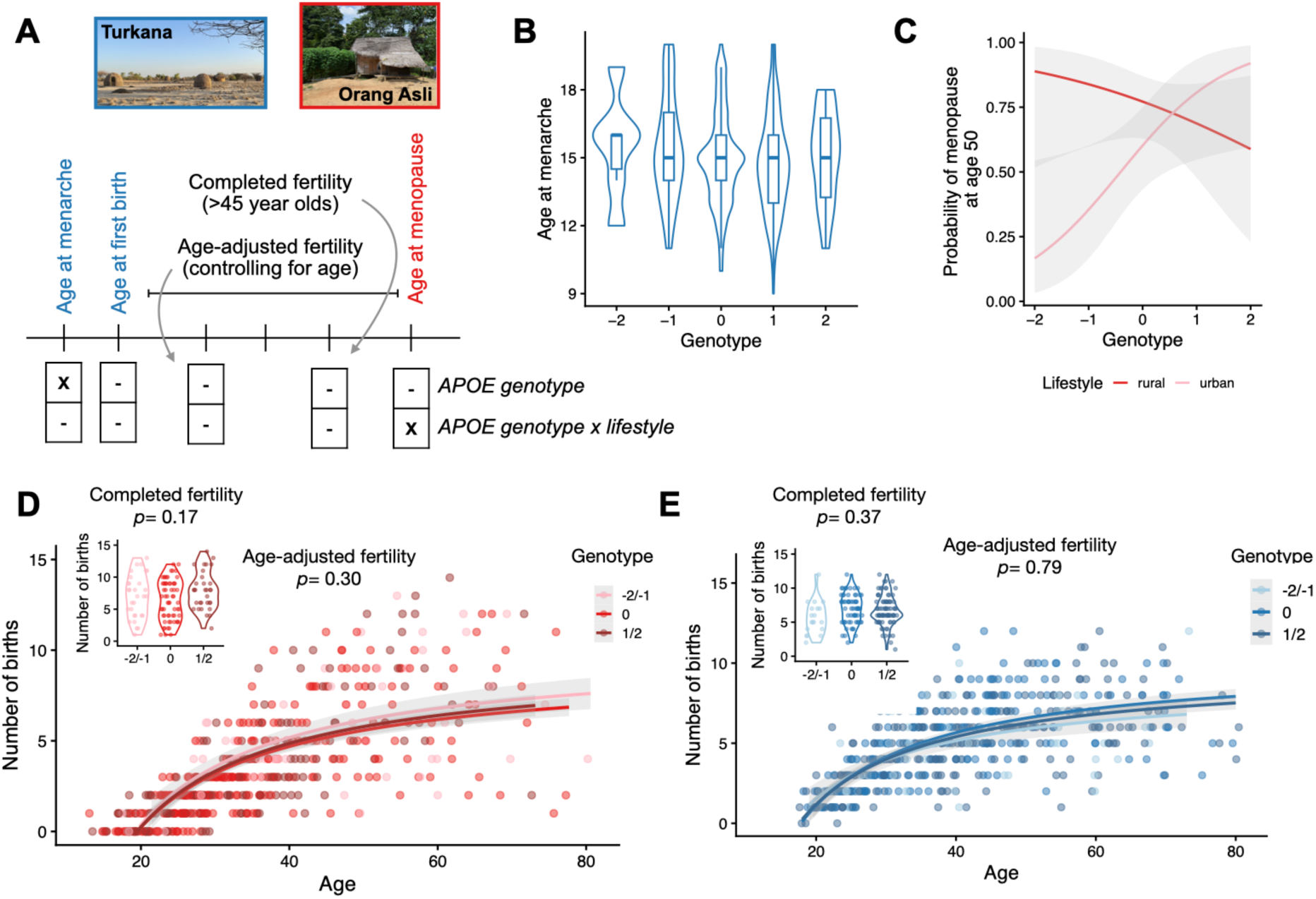
Select reproductive traits exhibit APOE genotype effects and genotype by lifestyle interactions. (A) Outline of key reproductive traits tested in each population with Turkana shown in blue and Orang Asli shown in red. Traits tested in both populations are labeled in black. The table below shows which traits exhibited significant effects, with the variables noted in rows. (B) Turkana ε4 carriers report younger average age at menarche. Box/violin plots generated from raw data. (C) Probability of reporting menopause at the age of 50 among Orang Asli of various genotypes. Genotype by lifestyle effects are driven by environmental sensitivity in ε2 carriers. Lines represent model predictions, shading represents the 95% confidence interval, with individuals binned into rural versus urban (although urbanicity was always modeled as a continuous variable). (D) APOE genotype is not associated with age-adjusted fertility or completed fertility (inset plot) in either Orang Asli or (E) Turkana. Lines represent linear model predictions of number of births as a function of age, fit separately for each APOE genotype group. Shading represents the 95% confidence interval and points represent the raw data.

## Discussion

Our work tests the hypothesis that the APOE ε4 isoform—which increases disease risk in older adults in urban, industrialized environments—has been maintained because it conferred benefits in the high-pathogen, energy-limited environments that characterized much of human evolution. While previous work has examined APOE genotype effects in non-industrial societies and shown differential patterns relative to those reported in urban, industrialized settings [15,22–24], our design leverages data from populations spanning non-industrial to industrialized contexts to directly test for moderating effects of lifestyle. We predicted that 1) APOE genotype would have consistent effects on cholesterol traits across lifestyles, but that 2) APOE immune trait correlations would be moderated by lifestyle. We also predicted that 3) APOE genotype-fertility associations would be stronger in non-industrial relative to industrialized settings (Figure 1).

Consistent with our first prediction, APOE ε4 carriers exhibited higher LDL and total cholesterol and lower HDL levels than APOE ε3 carriers, with no evidence for GxE interactions. Notably, APOE ε2 displayed mirror opposite effects on cholesterol traits—both in direction and magnitude—to those of APOE ε4. While this is not surprising given the known biochemical differences between these isoforms [2], previous work in non-industrial contexts has not always been able to evaluate APOE ε2 (e.g., due to its low frequency in South America) [11]. Previous work has shown that APOE ε2 is protective against Alzheimer’s and cardiovascular disease (relative to APOE ε3) in industrialized settings; however it also increases risk for several vascular and musculoskeletal diseases, suggesting complex costs and benefits [30]. Thus, evaluating the adaptive significance of APOE ε2 in environments relevant to human evolution is an important gap we aimed to fill. Because the Turkana and Orang Asli contain all three isoforms, we were able to construct a linear genotype variable that acknowledges and utilizes the differential impacts of APOE ε4 and APOE ε2.

APOE isoforms influence traits beyond cholesterol metabolism, with increasingly appreciated effects on innate immune function [2,12,57]. Consistent with the literature from industrialized settings [30,31], we found that APOE ε4 was generally associated with downregulation of innate immune activity in a baseline state (e.g., lower CRP levels, downregulation of innate immune genes). Consistent with our second prediction, we also found trends toward GxE interactions in the Orang Asli for multiple biomarkers: more APOE ε4 alleles trended toward associations with higher innate immune activation in rural, non-industrial contexts, but lower biomarker levels in urban, industrialized settings (i.e., urban individuals recapitulated patterns seen in industrialized contexts [30,31]). While suggestive, we cannot evaluate from these findings whether APOE ε4 is necessarily beneficial to immune function in non-industrial contexts. Doing so would require additional work connecting biomarkers to outcomes like pathogen clearance. However, our results point toward APOE ε4’s immunosuppressive phenotype being amplified by industrialization, and more generally to the idea that APOE ε4’s effects are environmentally flexible—an observation relevant to the puzzle of its persistence and global patterns.

Our findings on GxE interactions raise the following key question: what components of industrialized environments are important for modifying genotype-innate immune relationships? Previous work has suggested that sterile sources of damage-associated molecular patterns (DAMPs)—including obesity, inflammatory diets, smoking, or low physical activity levels—could be relevant perturbations to innate immune and inflammatory activity [15,39–43]. Our analyses of BMI, reliance on market-derived foods, tobacco usage, and objectively measured physical activity levels point towards excess adiposity and low activity levels as relevant to genotype x industrialization interactions. We note that our composite urbanicity index showed stronger associations with immune biomarkers than any single component, indicating that cumulative DAMP exposure in urban environments likely drives GxE patterns. However, a key limitation of our study is a lack of data on pathogen exposure and load across the lifestyle gradient in both populations: specifically, the high cholesterol levels and rapidly engaged immune phenotypes associated with APOE ε4 are thought to be beneficial under the combination of energy limitation and high pathogen burden [16,22–25]. Thus, a major avenue for future work is to characterize pathogen load and identity alongside cholesterol, immune, and other outcomes for the same individuals.

Finally, contrary to our third prediction, we did not observe the same robust association between APOE ε4 carrier status and female fertility in rural, non-industrial contexts, as was previously reported in energy-limited, high-pathogen settings in Bolivia, Ghana, and Ecuador [22–24]. We did find evidence for earlier maturation among APOE ε4 carriers and GxE effects on age at menopause, but these effects were modest. Our inability to detect genotype effects on fertility could be due to noise in self-reported data, or power that is lower than previous work in non-industrial groups (namely the Tsimane of Bolivia). Specifically, in our analyses of completed fertility (n = 122 and 165 in Orang Asli and Turkana, respectively), we estimate that we had <30% power to detect differences of 0.5 children between APOE ε4 carriers and non-carriers (e.g., the effect size reported for the Tsimane in [22]) (Figure 6D-E, Figure S7). While we were able to compare fertility outcomes among females who reported not currently using contraceptives, we did not have information about lifetime contraceptive usage. We also note that APOE genotype may have diverse effects on fertility correlates that we did not explore.

Genotype effects on hormones or immune tolerance could impact conception probabilities: higher luteal progesterone in APOE ε4 carriers may indicate higher fertility potential [58], while lower baseline inflammation in APOE carriers may facilitate implantation [59] (though associations between APOE ε4 and recurrent pregnancy loss are highly mixed [60]). Potentially most relevant, it is unclear how high the pathogen load is in Turkana and Orang Asli relative to the Tsimane, and differences in this key proximate determinant could thus drive variable fertility effects. Following birth, APOE genotype could impact infant or child mortality, with previous work showing that APOE ε4 carriers are less susceptible to the negative effects of enteric diseases during childhood [41,42]. Thus, while the literature has pointed toward APOE ε4 carriers producing more children in non-industrial, natural fertility settings, more work is needed to uncover the precise biological path to this outcome.

Our study has several limitations, most notably: 1) a lack of profiling of a proximate variable that is likely highly relevant for explaining GxE interactions (i.e., pathogen load and identity) and 2) a lack of data on disease outcomes relevant to APOE (e.g., cardiovascular disease and Alzheimer’s disease). This leaves clear directions for future work. Going forward, we are also excited for studies that move beyond baseline cell states to test the hypothesis that APOE ε4 carriers exhibit heightened responsiveness and pathogen clearance capacity [31,61] despite having lower baseline inflammation. For example, collecting transcriptomic data from individuals in both non-industrial and industrial settings, combined with ex vivo pathogen stimulations [62], would allow researchers to test whether GxE interactions shape the infection response. Such data could also be used to ask whether GxE interactions on innate immune function amplify at older ages as chronic, systemic inflammation increases, particularly among urban populations. Clarifying these dynamics will not only illuminate the evolutionary persistence of APOE variation but also refine our understanding of its modern health implications.

Our results add to growing evidence that APOE ε4, long considered a “detrimental” allele in industrialized settings, exerts pleiotropic and context-dependent effects that may have contributed to its maintenance. The widespread view of APOE ε4 as a deleterious allele overlooks its adaptive potential in pathogen-rich, energy-limited environments, where elevated lipid levels could have fueled immune responses, reduced infection-related mortality, facilitated faster growth, and maybe even provided fertility benefits (as shown in previous work but not here) [15,22–24]. Understanding these GxE interactions is important for evolutionary inference but also for informing personalized interventions [63]. For instance, APOE ε4 carriers may benefit from strategies that minimize chronic inflammatory burden (e.g., prompt management of infections, low-fat diets, regular physical activity) while avoiding unnecessary lipid suppression that could compromise immune function [17]. More broadly, these findings illustrate how evolutionary perspectives can clarify apparent health “paradoxes” by situating modern disease risk within the ecological and energetic contexts in which human genotypes evolved.

## Materials and methods

### Data collection and survey overview

We used data collected from two long-term, ongoing, integrated anthropological and biomedical studies: the Orang Asli Health and Lifeways Project (OA HeLP) and the Turkana Health and Genomics Project (THGP). OA HeLP data were collected between March 2022 and October 2024 and THGP data were collected between March 2018 and November 2023. For both Turkana and Orang Asli participants, informed consent was collected at multiple levels: first by first describing the project to the community as a whole and seeking the permission of community leaders, and subsequently through individual-specific review of the protocol and formal written consent. Only adults (18 years and older) were included in the study. Both studies engage communities across a wide lifestyle gradient and, relevant to this study, collect information on lifestyle and reproduction (through structured surveys) combined with biospecimens and health measurements (as described in [46,50]). Specifically, our lifestyle measurements relied on questions about engagement in wage labor as well as access to mobile phones, televisions, education, and toilets/sewage (as described in [47] and the Supplementary Materials). Female Turkana participants were also asked whether they were currently pregnant, whether they were on birth control, the number of children they had given birth to, and the age at which they had their first menstrual period and first child. Among female Orang Asli participants, interviewers asked participants whether they were currently pregnant, how many children and live births they had had, whether they were currently on birth control, and whether they had undergone menopause.

The OA HeLP was approved by the Medical Review and Ethics Committee of the Malaysian Ministry of Health (protocol ID: NMRR-20-2214-55565), the Malaysian Department of Orang Asli Development (permit ID: JAKOA.PP.30.052 JLD 21), and the Institutional Review Board of Vanderbilt University (protocol ID: 212175). The THGP was approved by Vanderbilt University (protocol ID: 00000162) and Kenya Medical Research Institute (KEMRI/SERU/CTMDR/119/4875).

### APOE genotyping

To identify the APOE isoform(s) that each individual carries, we extracted DNA from whole blood samples. For both THGP and OA HeLP, whole blood was collected in EDTA, immediately stored at –20°C, exported on dry ice, and subsequently stored at –80°C. DNA was extracted using the Quick-DNA Miniprep Plus or the Quick DNA/RNA extraction kits from Zymo Research and quantified using a Qubit Fluorometer. An aliquot of DNA was then shipped on dry ice to Johns Hopkins Nucleic Acid Technologies (JH-NAT) where samples were genotyped for the APOE ε2, ε3, and ε4 alleles using the TaqMan Allelic Discrimination system (Thermo Fisher Scientific, Carlsbad, CA). Allele determination was based on two SNPs (rs429358 and rs7412). In cases where either SNP was not confidently assigned, individuals were removed from further analyses. We tested whether rs429358 and rs7412 were in Hardy-Weinberg equilibrium using the HWExact function from the HardyWeinberg package in R [64].

### Cholesterol and innate immune biomarker measurements

For all participants, EDTA-collected whole blood samples were used to measure HDL, LDL, and total cholesterol levels with a CardioCheck Plus (PTS Diagnostics), and to conduct a 5-part white blood cell differential (with a HemoCue WBC Diff System). These analyses were done immediately after blood collection so they could be available in real time and used to inform primary health care provided alongside the research (as described in [50]). In our analyses, we focused on the following measures from the 5-part differential relevant to innate immune function: neutrophils (percent), lymphocytes (percent), monocytes (percent), neutrophils (total count), lymphocytes (total count), monocytes (total count), WBC (total count), neutrophil to lymphocyte ratio.

For a subset of Turkana participants, we also collected venous blood into a BD serum separator tube, spun the sample according to the manufacturer’s instructions, froze the sample at –20°C, and used the isolated serum to later measure CRP levels via a local provider (Columbia Africa in Nairobi, Kenya) using Turbidimetric methods. For a subset of Orang Asli participants, we also collected venous blood into a BD Cell Preparation Tube (CPT), spun the sample according to the manufacturer’s instructions, collected the plasma layer, and froze the plasma at –20°C. We later used these samples to measure APOE protein levels (via Targeted PRM Protein Analysis by LC-MS/MS) as well as cytokine and CRP levels (via the MSD VPLEX Human Proinflammatory Panel and CRP assay as described in [65]). These measurements were performed using aliquots sent to the Duke Proteomics and Metabolomics Core Facility and the Duke Molecular Physiology Institute Biomarkers Core Facility, respectively. The full list of measured cytokines was: TNF-a, IFN-γ, IL-10, IL-1B, IL-6, IL-12, IL-2, IL-4, and IL-8; we focus specifically on innate immune cytokines in the main text given our overall hypotheses. Finally, for a subset of Orang Asli participants, we also preserved the peripheral blood mononuclear cells isolated from the CPT tube for RNA extraction and transcriptomic analysis via mRNA-seq, following the methods detailed in the Supplementary Materials.

### Statistical analyses

We performed three main sets of statistical analyses to test the predictions that: 1) APOE ε4 would predict higher total and LDL cholesterol levels, and industrialized lifestyles would alter 2) APOE genotype-innate immune trait association as well as 3) APOE genotype-fertility trait associations. For all figures and summary statistics, if a tested APOE x lifestyle interaction effect was not significant (p > 0.05), we report estimates from an additive model that does not include the interaction. Wherever we tested genotype, lifestyle, or interactive effects on multiple grouped outcomes, we corrected for multiple hypothesis testing using a Benjamini-Hochberg FDR from the p-adjust function in R (R Core Team, 2023). We use the notation p_unadjusted_ to identify uncorrected p-values and p_FDR_ to identify FDR-adjusted p-values. In cases where no FDR correction was needed (e.g., because a single model was run to test a given biological hypothesis), we simply use the notation p.

#### Cholesterol outcomes

We asked whether APOE genotype and urbanicity predicted blood lipid levels in the Turkana and Orang Asli, as well as serum APOE protein levels in Orang Asli. For the analyses presented in the main text, we modeled APOE as a continuous variable, setting genotype ε4/ε4 to 2, ε3/ε4 to 1, ε3/ε3 and ε2/ε4 to 0, ε3/ε2 to –1, and ε2/ε2 to –2. Using linear models, we tested whether each outcome variable was predicted by age, sex, continuous APOE genotype, and urbanicity score. All variables except sex were centered and scaled.

#### Innate immune outcomes

We used a similar linear modeling framework to ask whether the following biomarkers of the innate immune system differed across lifestyles, APOE genotypes, and their interaction (controlling for age and sex): total white blood cell count, counts and percentage of neutrophils, monocytes, and lymphocytes, and CRP in Turkana and Orang Asli, as well as innate immune cytokine and PBMC gene expression levels in Orang Asli (information on the transcriptomic analyses is provided in the Supplementary Materials). Measures of C-reactive protein, blood cell counts, and cytokines were log₂-transformed in both groups to reduce skewness in their distributions. After identifying several GxE interactions for innate immune outcomes in the Orang Asli, we performed follow up analyses to assess whether the following proximate factors could recapitulate patterns seen in models using the urbanicity score: 1) body mass index (BMI) calculated as weight (kg) / height (m)^2^, 2) an index of how much access each individual has to sugar, salt, and oil (coded as 0=none, 1=some, 2=daily and summed across all 3 items) [47], 3) usage of tobacco products (coded as 0=never versus 1=sometimes or daily), and 4) objectively measured physical activity levels from wrist worn accelerometry (operationalized as mean Euclidean Norm Minus One or ENMO, derived from the GGIR package in R [66]) [52] (see Supplementary Materials). These analyses followed the same linear modeling strategy, swapping urbanicity for the variables described above. Finally, for comparison to previous work in the Tsimane [15], we tested whether lifestyle modified lipid effects on innate immune outcomes, focusing on the impacts of HDL, LDL, and cholesterol levels on the innate immune biomarkers measured in both populations (with all pairwise combinations of lipids and innate immune biomarkers explored). Here, we used linear models controlling for age, sex, main effects of the lipid of interest, main effects of urbanicity, and a lipid x urbanicity interaction.

#### Fertility outcomes

We filtered both datasets for a handful of values that were improbable or not internally consistent across outcomes (e.g., individuals that reported giving birth before reaching menarche; Supplementary Materials). We used linear and generalized linear models to predict fertility traits in both groups. Specifically, we used linear models including APOE genotype, urbanicity, and a genotype x urbanicity interaction to describe age at menarche (Turkana), age at first birth (Turkana), and age at menopause (Orang Asli). We used the same predictor variables and a generalized linear modeling approach with a Poisson error structure to describe completed fertility in females 45 years or older in both populations (following [22]). We also modeled number of births in all individuals using a generalized linear model with a Poisson error structure, but including an inverse age term (1/age) as well as APOE genotype, urbanicity, and a genotype x urbanicity interaction.

#### Genetic relatedness and population structure

A common problem in genotype-phenotype association studies is improper control for genetic relatedness and population structure [67,68]. In other words, false positive results can occur because related individuals are more likely to have similar genotypes as well as similar phenotypes, even if genotype does not affect phenotype. To the best of our knowledge, the THGP dataset contains minimal close relatives (e.g., a single individual per household is invited to participate); this aspect of sampling design, as well as a general absence of confounding between lifestyle and population structure, has previously been tested for and described [69]. For Orang Asli, certain ethnolinguistic groups are more likely to reside in rural versus urban areas, and there is some subtle genetic differentiation between subgroups [70,71]. Therefore, we repeated all of our main analyses in Orang Asli using linear mixed effects models implemented in the R package EMMREML [72]. These models incorporate a random effect parameterized by a genetic relatedness matrix, which captures both close kinship and broader population structure [67]. Pairwise genetic relationships were inferred from genotypes derived from the transcriptomic data (see Supplementary Materials) and estimated in Plink [73]. These supplementary analyses agreed with the results provided in the main text and are detailed in Table S17.

We carried out all statistical analyses in the R version 4.4.2 environment.

## Data and code availability

The OA HeLP and THGP prioritize minimizing risk to study participants and follow the CARE Principles for Indigenous Data Governance (emphasizing Collective Benefit, Authority to Control, Responsibility, and Ethics) and the FAIR Guiding Principles for scientific data (emphasizing Findability, Accessibility, Interoperability, and Reusability). Individual-level data are stored in a public but protected repository at Zenodo (10.5281/zenodo.18686419) and available through restricted access. Requests for de-identified data must include a detailed application and procedures for data security, privacy, and minimizing potential harm. Sample data use agreements are provided at https://lea-lab.org/resources.html.

Code to perform all analyses described here is available on GitHub as follows: 1) generating the urbanicity score for both populations (https://github.com/mwatowich/Multi-population_lifestyle_scales); 2) performing statistical analyses and processing the accelerometry data (https://github.com/mwatowich/effects_of_APOE_across_industrialization_gradient); 3) processing the Orang Asli transcriptomic data (https://github.com/laylabrassington/ELvsCL/blob/main/scripts/OA_RNAseq_data_processing.R md).

## Acknowledgments

First and foremost, we thank Turkana and Orang Asli community members and leaders for their contributions, hospitality, and support of our scientific work. We also thank all present and previous members of the Turkana Health and Genomics Project and the Orang Asli Health and Lifeways Project. We are grateful to the staff of the Kenya Medical Research Institute for their essential support. The Hospital Orang Asli Gombak, Malaysian Red Crescent Society, Federation of Private Medical Practitioners’ Associations of Malaysia, and Center for Orang Asli Concerns provided critical support for data collection for the Orang Asli Health and Lifeways Project.

