## Supplemental Methods and Figures for "Divergent Biological Consequences of APOE Isoforms Across Industrialized and Non-Industrial Environments"

**This file includes:**


- Supplementary Materials and Methods
- Figure S1. Maps of participant villages across a lifestyle gradient in Kenya and Malaysia
- Figure S2. Opposing effects of APOE ε4 and APOE ε2 isoforms on cholesterol
- Figure S3. Genotype, lifestyle, and GxE interaction effects on innate immune biomarkers
- Figure S4. Genotype effects on Orang Asli immune gene expression levels
- Figure S5. Cholesterol, lifestyle, and cholesterol x lifestyle effects on innate immune biomarkers
- Figure S6. Using different age cutoffs to define completed fertility does not impact conclusions
- Figure S7. Power to detect genotype effects on fertility
- Supplementary References


**The following are provided in a separate file:**


- Table S1. Orang Asli participants broken down by self reported ethnolinguistic group
- Table S2. Turkana and Orang Asli participant demographic information
- Table S3. Results from linear models testing for lifestyle effects on body fat percentage and body mass index
- Table S4. Results from Haldane Exact tests for Hardy-Weinberg equilibrium
- Table S5. Results from linear models testing for genotype (number of ε4 or ε2 alleles) and urbanicity effects on cholesterol measurements
- Table S6. Results from linear models testing for genotype (coded linearly) and urbanicity effects on cholesterol measurements
- Table S7. Percent variance explained by linear model covariates testing for genotype and urbanicity effects on cholesterol measurements
- Table S8. Results from linear models testing for genotype (coded linearly) and urbanicity effects on immune biomarkers in Orang Asli
- Table S9. Results from linear models testing for genotype (number of ε4 allele) and urbanicity effects on immune biomarkers in Orang Asli
- Table S10. Results from linear models testing for genotype (coded linearly) and urbanicity effects on immune biomarkers in Turkana
- Table S11. Results from linear models testing for genotype (number of ε4 allele) and urbanicity effects on immune biomarkers in Turkana
- Table S12. Results from linear models testing for effects of genotype and proximate lifestyle variables on immune biomarkers in Orang Asli

- Table S13. Results from linear models testing for cholesterol and urbanicity effects on immune biomarkers in Orang Asli
- Table S14. Results from linear models testing for cholesterol and urbanicity effects on immune biomarkers in Turkana
- Table S15. Results from linear models testing for genotype and urbanicity effects on fertility traits in Orang Asli
- Table S16. Results from linear models testing for genotype and urbanicity effects on fertility traits in Turkana
- Table S17. Main results (Orang Asli data only) recapitulated using linear mixed effects models

### Supplementary Materials and Methods

#### The Turkana of Northwest Kenya

The Turkana are historically a nomadic pastoralist population living in the remote Turkana Basin in northwest Kenya. Traditionally, high amounts of the Turkana diet has been derived from animal products including milk, meat, and blood, though these numbers vary highly depending on the season and geographic area [1,2]. Ongoing infrastructure construction and rapid economic development of Kenya has resulted in the growth of several urban centers in and near traditional Turkana lands, the expansion of small-scale markets, and an increased reliance on agriculture. As a result, most Turkana no longer exclusively practice traditional pastoralism, instead relying on trade, small scale farming, and increasing participation in the market economy. In addition to socioeconomic changes happening within the Turkana region, many Turkana have moved to highly urbanized areas in central Kenya in the last several decades, and now participate fully in the market-economy [1,3].

We used data collected by the Turkana Health and Genomics Project (THGP) (March 2018-November 2023) described in Lea et al [1]. Following meetings with community leaders and members, the THGP project recruited consenting, healthy adults (>18 years) to participate in structured surveys with a research team member who was familiar with the local community and language of the participant (e.g., Turkana, Swahili, English). All community members, regardless of whether they participated in the project, were provided free primary healthcare and consultation from a local nurse working with the project if desired. Individuals who wished to participate took part in structured interviews asking about demography, subsistence and labor practices, material wealth and housing infrastructure, diet, health, and other aspects of lifestyle. Written, informed consent was obtained from all participants after the study goals, sampling procedures, and potential risks were explained to participants in their native language by researchers. Data collection is further described in [1,4].

#### The Orang Asli of Peninsular Malaysia

“Orang Asli” refers broadly to the Indigenous peoples of Peninsular Malaysia. The Orang Asli are typically categorized into three broad groups (the Negrito or Semang, Senoi, and Aboriginal Malay) and 19 culturally distinct ethnolinguistic groups. While these three subgroups are somewhat genetically distinct, the Orang Asli are generally much more genetically similar to each other than to ethnic Malays or other Asian populations [5]. Collectively, the Orang Asli traditionally practiced varied subsistence histories consisting of foraging, swidden agriculture, trade of rainforest products, or a combination thereof, and these practices were ubiquitous among Orang Asli until the 1950’s and remain common in some areas today [6,7]. Over the last several decades, Malaysia has undergone one of the fastest rates of urbanization, deforestation, and industrialization worldwide [8,9], which, combined with government efforts to assimilate the Orang Asli into the national economy, has resulted in Orang Asli experiencing a wide gradient of lifestyles. This ranges from individuals living in remote interior rainforest villages and practicing small-scale, subsistence level lifestyles such as foraging and hunting to Orang Asli villages being surrounded by urban or peri-urban development and individuals participating in the market economy instead of, or in addition to, traditional practices [7].

We used data from the Orang Asli Health and Lifeways Project (OA HeLP) that were collected between March 2020 and October 2024. OA HeLP works in Orang Asli communities and performs similar sampling and data collection as the THGP. OA HeLP also provides free primary healthcare to community members regardless of participation status in collaboration with local hospitals and NGOs. After consent from community leaders and hosting community-wide informational sessions, OA HeLP invited adults to participate in the study. The study aims, procedures, and potential risks were explained to all participants and written, informed consent was obtained from all participants. Additional information about OA HeLP is described in [10].

#### Urbanicity scale generation

Turkana and Orang Asli live across wide lifestyle gradients, which we have previously described using a location-based "urbanicity" score [11–14]. Using a location-level scale is advantageous because it allows individuals who might be missing individual-level data to be included in analysis and because it gives a representation of the resources available across the community (e.g., an individual may not own a television but may have access through family or friends). To quantify individuals' exposure to urban centers and urban, industrialized, and market-integrated infrastructure comparably in both populations, we used a scale that was first proposed by [15]. This scale was tested in Orang Asli and Turkana in [11] and predicted cardiometabolic health better than other measures of urbanicity, acculturation, industrialization, and market-integration. Scale construction is shown below. Population density was estimated from NASA's Gridded Population of the World resource with a resolution of 2.5 arc-minutes [16]. The urbanicity score values ranged from 4 - 41.8 in Orang Asli and between 0.67 - 41.4 in Turkana, emphasizing highly comparable lifestyle gradients. We modeled urbanicity score as a continuous variable in our analyses but for the purposes of plotting, we show data from "rural" and "urban" individuals following a natural break in the bimodal distribution of urbanicity in both populations at 25.

| **Population density (estimated number of people per square kilometer)** | **Contribution to scale** |
|---|---|
| 0-100 | 1 |
| 100-200 | 2 |
| 200-300 | 3 |
| 300-400 | 4 |
| 400-500 | 5 |
| 500-1000 | 6 |
| 1000-2000 | 7 |
| 2000-3000 | 8 |
| 3000-4000 | 9 |
| 4000-6000 | 10 |

| | |
|---|---|
| 6000-8000 | 11 |
| 8000-10000 | 12 |
| 10000-15000 | 13 |
| 15000-20000 | 14 |
| >20000 | 15 |
| **Occupation** | |
| Proportion of the population involved in non-wage labor | 10 - (10 x (proportion)) |
| **Built Environment** | |
| Proportion of households with flush toilets | 5 x proportion |
| Proportion of households with electricity | 5 x proportion |
| **Communication/market-derived items** | |
| Proportion of households with mobile phone | 5 x proportion |
| Proportion of households with television | 5 x proportion |
| **Education** | |
| Proportion of surveyed individuals > 40 years with any level of formal education | 10 x proportion |
| Proportion of surveyed individuals < 40 years with any level of formal education | 10 x proportion |

#### Orang Asli accelerometry data collection and processing

OA HeLP study participants were given a triaxial accelerometer (Axivity AX3, Axivity Ltd., UK) to wear for up to 10 days on their non-dominant wrist to measure their physical activity patterns (median = 6 wear days per person). The devices were later collected by a community volunteer and eventually returned to a member of the research team. Accelerometry is purposefully restricted to periods when researchers are not working in the community to ensure that behavior is not altered by the presence of the research team.

Prior to deployment of the accelerometers, the devices were configured to collect data at a sampling frequency of 100 Hz with a dynamic range of 8 g. Raw accelerometry files were downloaded and pre-processed to remove any days recorded after participants wore the devices. We then used the GGIR package v. 3.2.6 in R v. 4.4.2 to process data into 5-second epochs for analysis using default auto-calibration and imputation algorithms [17,18]. Non-wear time was estimated using built-in GGIR algorithms, and we defined a valid day for inclusion as those in which ≥16 hours of wear time were detected. We extracted the Euclidean Norm Minus One metric—a common summary measure of overall activity levels for the day—from the GGIR output. Specifically, this metric is the Euclidean norm (vector magnitude) of the x, y, and z axes (for a tri-axial device) representing raw signals of acceleration, accounting for the effect of gravity by subtracting one gravitational unit.

#### Orang Asli gene expression data generation and processing

We measured gene expression levels in peripheral blood mononuclear cells isolated from whole blood for 755 Orang Asli individuals. To do so, approximately 8mL of venous blood was collected from each participant in a CPT tube, from which peripheral blood mononuclear cells (PBMCs) were isolated following the manufacturer's instructions. PBMCs were preserved in Zymo's DNA/RNA shield and stored in liquid nitrogen for transport followed by longer-term storage at -80C. RNA was extracted from the PBMCs using Zymo Quick-RNA 96 kits, Zymo Quick-DNA/RNA MagBead kits, or Zymo Quick-RNA MiniPrep Plus kits and were library prepped using the NEBNext Ultra II RNA library prep kit. Samples were molarity-normalized and sequenced on the Illumina NovaSeq X sequencing platform at The Translational Genomics Research Institute (TGen) or the Vanderbilt University Medical Center (VUMC) Vanderbilt Technologies for Advanced Genomics (VANTAGE) core. All samples were sequenced to an average depth of 18,142,612 total reads (range = 3,246,661 - 67,538,791).

We followed standard pipelines for read trimming and removing adapter contamination as described in [19–21]. Next, reads were mapped to the human reference genome (hg38) using STAR [20] and gene counts were compiled using HTSeq [21]. As further quality control, given that our PBMC samples were isolated from whole blood, we calculated the percent of reads mapping to hemoglobin-related genes (specifically HBA1, HBA2, and HBB) and removed any samples with >10% hemoglobin-associated reads. We also excluded any samples with low read counts (<1 million reads), low mapping rates (<70% of reads uniquely mapped to the human genome), or whose sex inferred from the transcriptomic data (i.e., from Y chromosome read counts) did not match their self reported sex (indicating a potential sample mix up or labeling issue).

#### Orang Asli genetic relatedness matrix construction

We performed a set of supplementary analyses in Orang Asli that controlled for pairwise genetic relatedness using linear mixed effects models. To generate these values, we first derived individual-level genotype calls from the mRNAseq reads. To do so, aligned BAM files were processed using a variant-calling pipeline based on GATK (v4.1.4.0) and SAMtools (v1.9) [22,23]. Briefly, coordinate-sorted BAMs were filtered to retain only autosomal reads and indexed with SAMtools. Duplicate reads were identified and marked using Picard (v2.18.27), and read group information was added to ensure compatibility with downstream GATK tools [24]. GATK's SplitNCigarReads was used to adjust spliced alignments for RNA-seq data. GATK's base quality score recalibration (BQSR) was performed using known high-confidence SNPs from the 1000 Genomes Project phase 1 as reference sites [25].

Variant calling was conducted per sample using GATK's HaplotypeCaller in gVCF mode, followed by variant quality filtering to remove calls with Fisher Strand (FS > 30.0) or low quality-by-depth (QD < 2.0). The resulting per-sample filtered VCFs were zipped, indexed, and merged across all individuals with bcftools (v1.18) [26]. Only variants with the PASS filter flag were retained for subsequent analyses. The merged multi-sample VCF was further filtered to remove variants with excessive missingness (allele number < 50% of samples), minor allele frequency (MAF) < 1%, or Hardy-Weinberg equilibrium ($p<1\times10^{-6}$), using PLINK (v1.9) [27]. After all filtering steps, high-quality biallelic SNPs were retained for genotype based quality control.

Pairwise kinship coefficients were computed using KING (v2.3.2) to confirm sample independence and identify potential duplicates [28]. The final high-confidence genotype matrix was used to generate a pairwise genetic relatedness matrix in Plink for downstream linear mixed effects modeling.

#### Orang Asli gene expression data analysis

To test whether gene expression levels differed across APOE genotypes, we used the R package EMMREML [29] to employ a linear mixed modeling approach. Gene expression counts were first filtered to remove lowly expressed genes (average transcripts per million > 2), normalized using the voomWithQualityWeights function in the R package limma [30], and corrected for sequencing batch using lmFit in limma. We modeled the normalized, batch-corrected expression values for 9993 genes using models that included the effects of APOE genotype (linear variable), while controlling for urbanicity, age, sex, percent of lymphocytes and monocytes from complete blood count with differential, and genetic relatedness between individuals. We corrected for multiple hypothesis testing using the qvalue package in R [31].

Though we observed a non-uniform distribution of genotype-associated p-values, indicating a likely global effect, we did not find any genes that were significantly associated with genotype at a 10% false discovery rate cutoff. Nevertheless, to identify biological processes that were enriched in genes with the largest APOE genotype effect sizes, we conducted a gene set enrichment analysis using the fgsea package in R [32], which preranks genes by their effect size and does not require a set of significantly differentially expressed genes as input.

#### Outlier detection and filtering of lipid, innate immune, and fertility datasets

Before modeling our cholesterol, innate immune biomarker, or fertility datasets, we first removed outliers. We did not observe any obvious outliers in the three blood lipids we measured (range of lipid values for Orang Asli: total cholesterol = 100-400, HDL = 20-120, LDL = 29-288; range of lipid values for Turkana: total cholesterol = 99-332, HDL = 19-121, LDL = 1-188). We performed a log2 transformation of immune cell counts, cytokine values, and C-reactive protein and removed values that surpassed an interquartile range (IQR) of 2.

For our menopause analyses, if women reported being menopausal but also reported that they were currently cycling, pregnant, or breast feeding we changed these responses to pre-menopausal. We also excluded observations where menopause occurred prior to the age of 40 or after 62, as menopause occurring outside of this age range may have a different etiology than typical menopause [33,34]. In a very large, US representative dataset (NHANES), age at menarche ranged from 8-20 years old and thus we removed entries with age at menarche outside this range in our data (n=7). Following [35], we also excluded observations where age at first birth was greater than 30, as this likely reflected extenuating reproductive circumstances. Lastly, we excluded observations where the number of births exceeded the number of pregnancies (n=1) or the number of reproductively mature years would not allow for the number of live births reported (n=3) as these were likely recording errors.

#### Power analyses

We estimated power to detect differences in completed fertility between APOE ε4 carriers and non-carriers using a simulation-based framework that mirrors the Poisson generalized linear model (GLM) approach used in the main text. First, we specified the observed proportion of individuals carrying at least one APOE ε4 allele in each population (35.2% in Orang Asli and 46.4% in Turkana) and set the total sample size to a number varying from 50 to 400 individuals. For each sample size, we simulated 1000 replicate datasets in which the number of children for non APOE ε4 carriers was generated from a Poisson distribution with a mean of 6.5 (approximately the average number of children in the completed fertility dataset for both groups). APOE ε4 carriers were then assigned an increased mean of 0.5 children, consistent with previously reported effect sizes [35]. Each simulated dataset was analyzed using a Poisson GLM with genotype as a predictor, and the proportion of simulations in which the genotype coefficient achieved $P < 0.05$ was taken as the estimated power for a given sample size. Power curves were generated separately for the Orang Asli and Turkana genotype frequencies to illustrate how population-specific allele frequencies influence detectable effect sizes.

#### Supplementary Figures

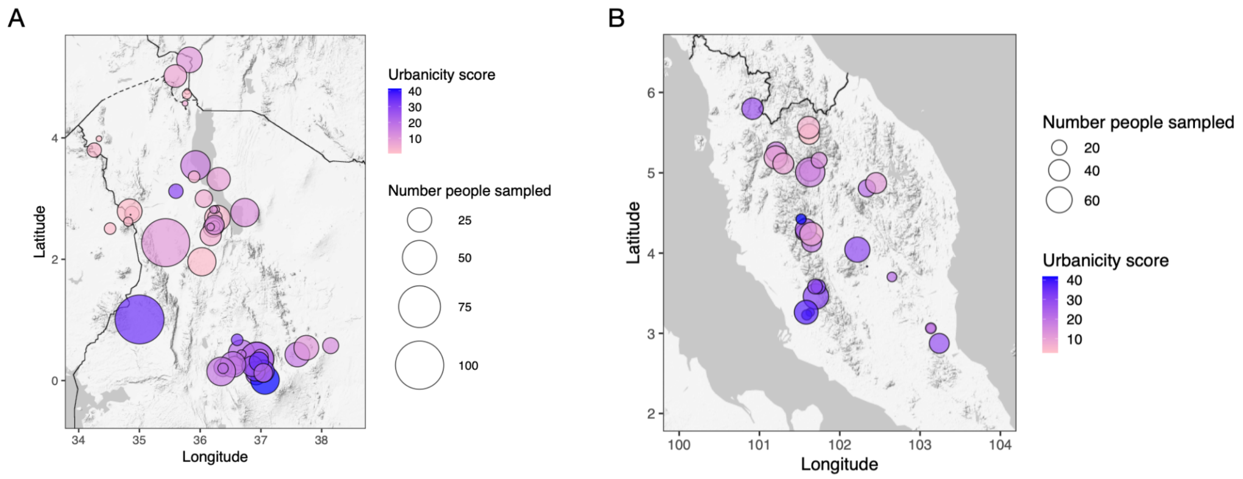


**Figure S1. Maps of participant villages across a lifestyle gradient in Kenya and Malaysia**. Panel A shows participant communities associated with the Turkana Health and Genomics Project (map of Kenya) and panel B shows participant communities associated with the Orang Asli Health and Lifeways Project (map of Peninsular Malaysia). Shading denotes the urbanicity score for each location and the size of the circle indicates the number of participants.

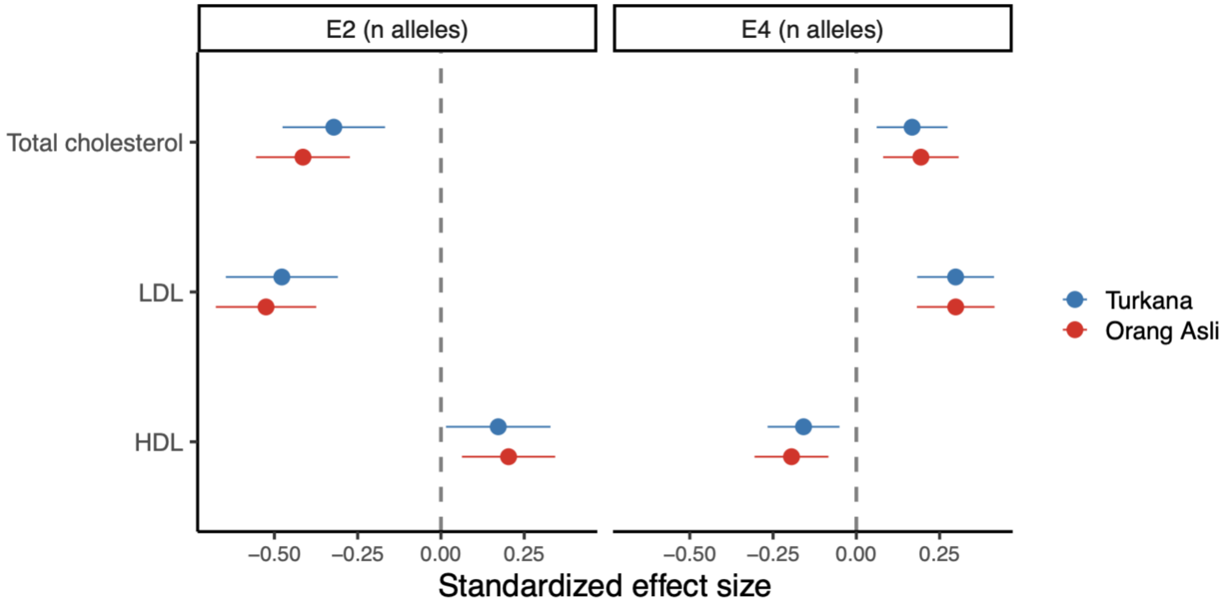


**Figure S2. Opposing effects of APOE ε4 and APOE ε2 isoforms on cholesterol.** Forest plot of the effect of APOE (coded as the number of APOE ε4 or APOE ε2 alleles) on cholesterol measures, controlling for age, sex, and urbanicity. Points represent the standardized effect size and bars represent 95% confidence intervals. Positive values indicate positive associations between cholesterol traits and allele number. All tests passed a 5% false discovery rate cutoff.

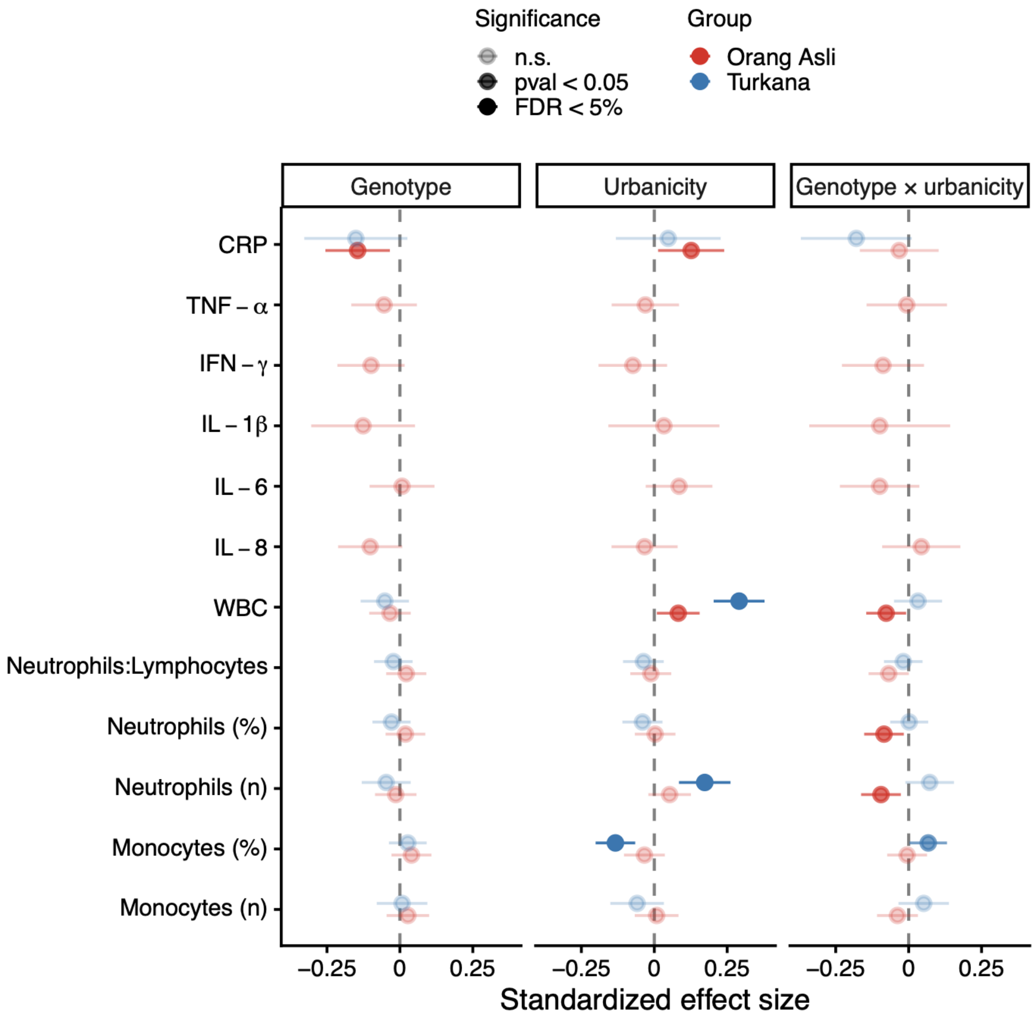


**Figure S3. Genotype, lifestyle, and GxE interaction effects on innate immune biomarkers.** Forest plot of effects of genotype (coded linearly), urbanicity, and genotype x urbanicity from a model including controlling for age and sex. Points represent the standardized effect size and bars represent 95% confidence intervals. Positive values indicate positive associations between immune traits and number of APOE ε4 alleles or increasing urbanicity.

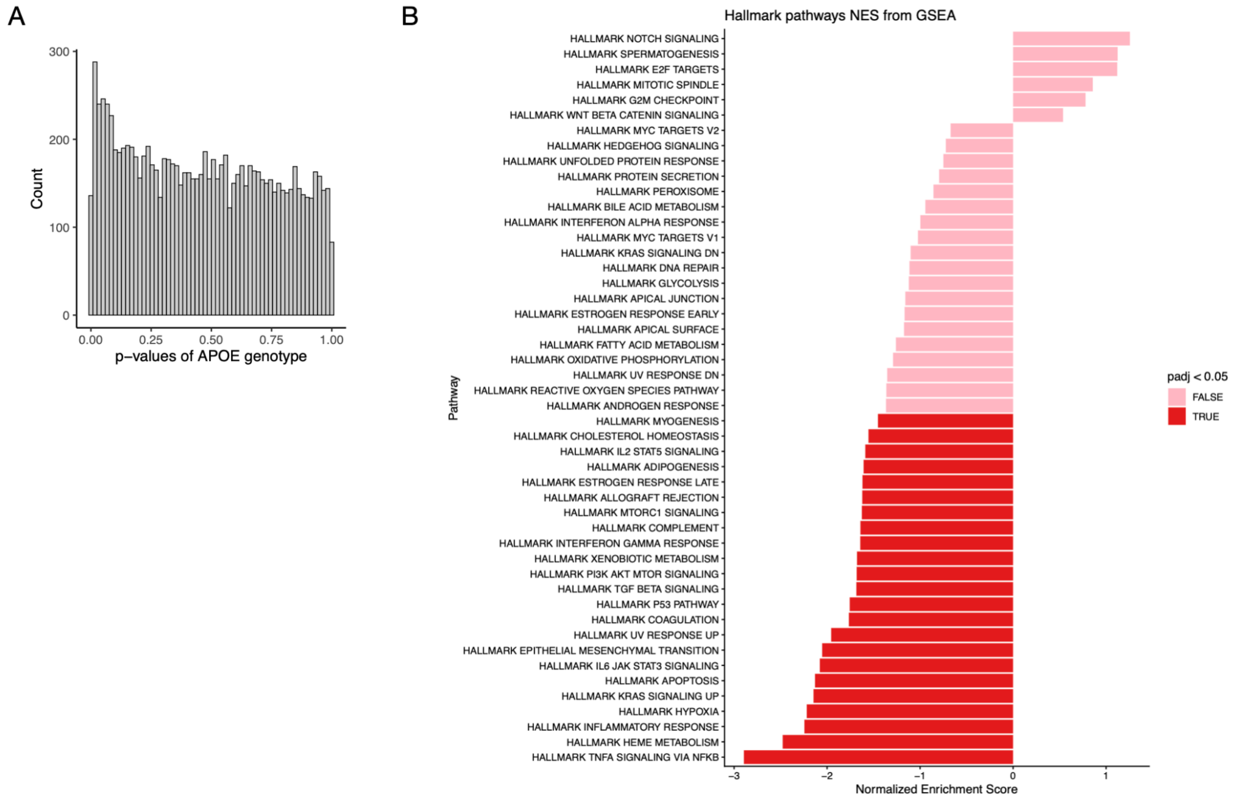


**Figure S4. Genotype effects on Orang Asli immune gene expression levels.** (A) Distribution of p-values for the linear APOE genotype variable, across all 9993 analyzed genes. P-values were extracted from linear mixed effects models controlling for age, sex, cell type composition, urbanicity, and genetic relatedness. (B) Gene set enrichment analysis scores, indicating degree of enrichment in a given pathway, for GSEA hallmark pathways. Pathways are colored by whether they passed a 5% false discovery rate threshold.

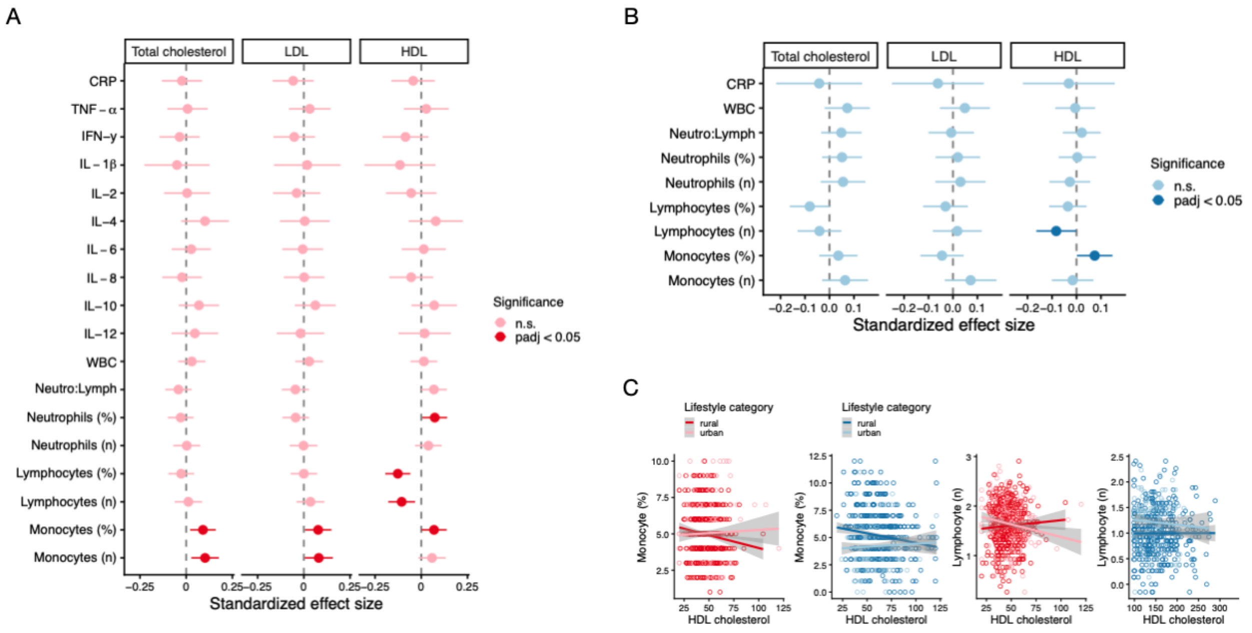


**Figure S5. Cholesterol, lifestyle, and cholesterol x lifestyle effects on innate immune biomarkers** in (A) Orang Asli and (B) Turkana. Forest plots show the magnitude of lifestyle x cholesterol trait interactions on each immune outcome from a linear model controlling for age and sex and corresponding main effects. Points represent the standardized effect size and bars represent 95% confidence intervals. (C) Shows example plots of relationships with significant interactive effects. Data from Orang Asli are shown in the first two plots in red and the third plot shown in blue is from the Turkana.

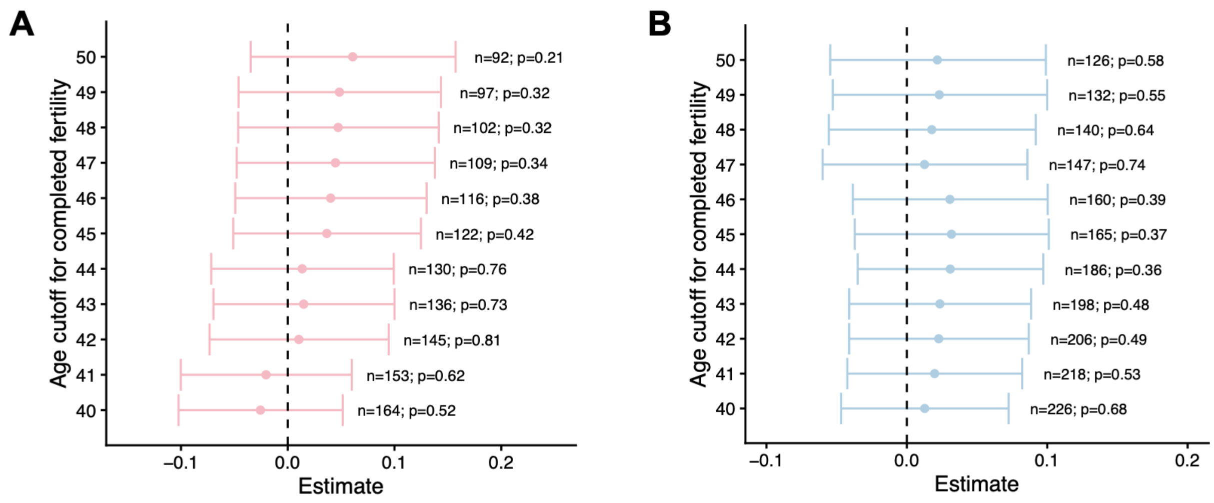


**Figure S6. Conclusions are robust to alternative age cutoffs used to define completed fertility.** Model estimates for the interaction between APOE genotype and urbanicity in predicting a woman's lifetime number of births in our completed fertility datasets from the Orang Asli (A) and Turkana (B). Across all age cutoffs considered, we found no significant interaction between APOE genotype and urbanicity. For each cutoff, sample sizes of women included in the completed fertility dataset and the corresponding p-values for the interaction term are reported.

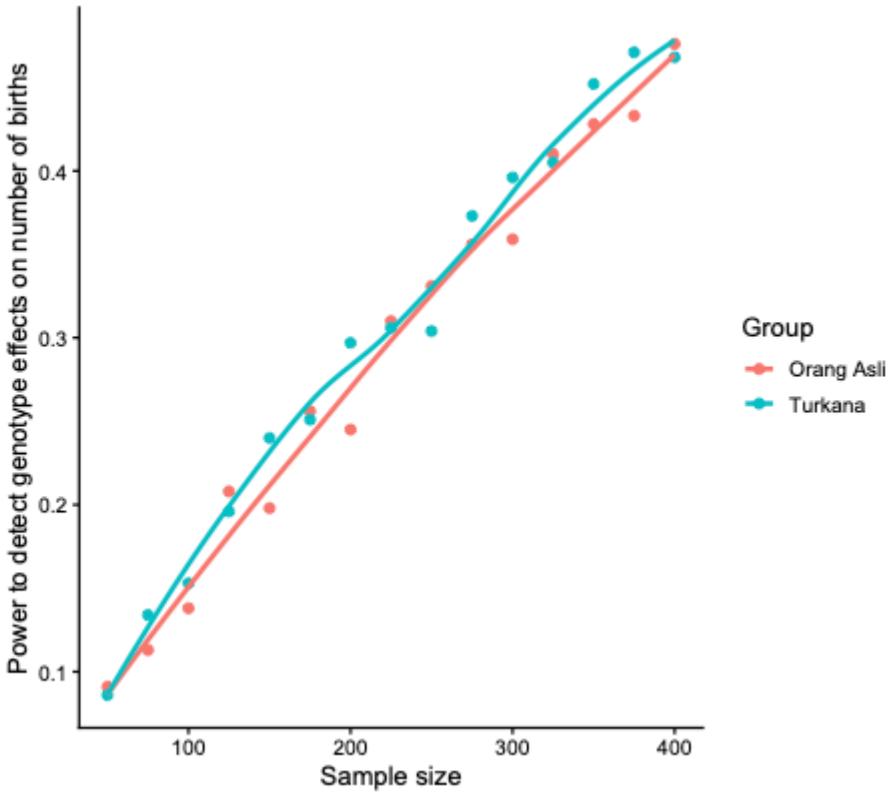


**Figure S7. Power to detect genotype effects on fertility.** We performed 1000 replicate simulations for datasets of varying sizes to estimate our power to detect a difference of 0.5 births between APOE ε4 carriers and non-carriers. Power was estimated as the proportion of simulated true effects detected at $p < 0.05$ in a generalized linear modeling framework. Power is generally low and has not asymptoted for our sample size range.
